# OPUS-BFactor: Predicting protein B-factor with sequence and structure information

**DOI:** 10.1101/2024.07.17.604018

**Authors:** Gang Xu, Yulu Yang, Ying Lv, Zhenwei Luo, Qinghua Wang, Jianpeng Ma

## Abstract

Protein B-factor, also known as the Debye-Waller factor, measures the fluctuation of an atom around its average position. It serves as a crucial indicator of protein flexibility and dynamics. However, accurately predicting the B-factor of C_α_ atoms remains challenging. In this work, we introduce OPUS-BFactor, a tool for predicting the normalized protein B-factor. OPUS-BFactor operates in two modes: the first mode, OPUS-BFactor-seq, uses sequence information as input, allowing predictions based solely on protein sequence; the second mode, OPUS-BFactor-struct, uses structural information, requiring the 3D structure of the target protein. Evaluation on three test sets, including recently released targets from CAMEO and CASP15, demonstrates that OPUS-BFactor significantly outperforms other B-factor prediction methods. Therefore, OPUS-BFactor is a valuable tool for predicting protein properties related to the B-factor, such as flexibility, thermal stability, and region activity.

## Introduction

Protein B-factor, also known as the Debye-Waller factor or temperature factor, measures the mean squared displacement or uncertainty of atoms. It describes the attenuation of X-ray or neutron scattering due to thermal motion in protein crystallography ^1, 2^. Numerous studies have shown that protein B-factor is valuable in various areas, such as predicting protein flexibility ^3, 4^, evaluating thermal stability ^5^, analyzing active and disordered regions ^6, 7^, and studying protein dynamics ^8^. Since protein fluctuation provides a crucial link between structure and function ^1, 9^, accurately predicting protein B-factor is essential for understanding the characteristics of target proteins.

Over the past several decades, numerous methods have been proposed for predicting protein B-factor ^1, 10-14^. In the early stages, most researches focused on developing physics-based models that require structural information of target protein. Pang sampled atomic positional fluctuations in multi-picosecond molecular dynamics (MD) simulations to predict crystallographic B-factors of folded globular proteins ^13^. In addition, some researches introduced normal mode analysis (NMA) into this field ^9^. In NMA, the Hessian of the harmonic potential is employed to describe the atomic thermal fluctuations, therefore, the B-factors of proteins are correlated with the eigenvalues of the Hessian. Lu *et al*. developed an all-atom normal mode calculation method, fSUB, and demonstrated its ability to model all-atom B-factor using normal-modes ^15^. Poon *et al*. utilized normal modes to fit the B-factor in crystallographic refinement and improve the accuracy of structure determination ^16^. Meanwhile, the Gaussian network model (GNM) and the anisotropic network model (ANM) are two elastic network models that have been widely used to study protein fluctuation dynamics ^17, 18^. They simplify the complicated all-atomic potential to a quadratic function near the equilibrium state, allowing the decomposion of motions into normal modes with different frequencies ^19^.

In recent years, several machine learning-based models have been proposed for predicting protein B-factors ^1, 10, 11, 20, 21^. Some of these models utilize support vector regression (SVR) ^10, 11, 20^, and some of them employ graph models, such as multiscale weighted colored graphs ^21^. With the development of deep learning techniques ^22^, several new methods based on deep learning frameworks have emerged ^12, 14^. Most of these methods adopte the Bidirectional Long Short-Term Memory (BiLSTM) network ^23^, which largely improve the accuracy of B-factor prediction.

In this study, we introduce a deep learning-based model named OPUS-BFactor for predicting protein B-factors (specifically for C_α_ atoms). OPUS-BFactor operates in two modes. In the first mode, it uses sequence information as input, enabling predictions based solely on protein sequence. Previous sequence-based methods typically rely on one-hot encoding ^12^ or evolutionary features such as PSSM profiles (Position Specific Scoring Matrix) and HHM profiles (Hidden Markov Model) ^14^. In this study, OPUS-BFactor adopts a more powerful evolutionary feature derived from the protein language model ESM-2 ^24^. The results show that using ESM-2 features as inputs significantly improves B-factor prediction accuracy compared to those using one-hot encoding and PSSM features. In the second mode, OPUS-BFactor utilizes structural information, achieving better results than the sequence-based mode. For clarity, we refer to the results of OPUS-BFactor based on sequence information as OPUS-BFactor-seq (first mode) and the results based on structural information as OPUS-BFactor-struct (second mode).

We assess the performance of OPUS-BFactor using three test sets: CAMEO65, CASP15, and CAMEO82. The results indicate that OPUS-BFactor-struct significantly outperforms other methods. Specifically, on the most recently released CAMEO82 test set, the average Pearson correlation coefficient (PCC) for B-factor from C_α_ atoms is 0.67 for OPUS-BFactor-struct and 0.58 for OPUS-BFactor-seq, compared to 0.41 for the most recent method proposed by Pandey *et al.* ^12^.

Although many methods have been proposed for protein B-factor prediction, they usually use different training and test sets, making fair comparisons difficult. Additionally, the code for many methods is not publicly available, complicating their use by other researchers. Therefore, we will make our training and test sets, as well as our code, available to all researchers. We hope that OPUS-BFactor will serve as a fair baseline method in protein B-factor prediction. Additionally, the formatted datasets may become a useful benchmark to facilitate the development of protein language models, given that the performance of sequence-based B-factor prediction models still lags behind that of structure-based models.

## Method

### Framework of OPUS-BFactor

OPUS-BFactor adopts the RotaFormer module from OPUS-Rota5 ^25^ as its backbone architecture, with some modifications. As shown in Figure 1, the 1D and 2D features are derived from protein structural information, while the ESM-2 features are obtained from the protein language model ESM-2 ^24^, which relies solely on protein sequence. Specifically, the 1D protein features include two one-hot encoded features for the 3-state and 8-state secondary structures, seven physicochemical properties ^26, 27^, 19 PSP features representing 19 rigid-body blocks within residues ^27-29^, and six backbone torsion angle features (sine and cosine values for □, ψ, and ω). The 2D features describe residue-residue backbone contact information ^30, 31^, including C_β_- C_β_ distance distributions and orientational distributions of three dihedrals (ω, θ_ab_, θ_ba_) and two angles (φ_ab_, φ_ba_) between residues a and b. Distances of C_β_- C_β_ span from 2 to 20 Å, segmented into 36 bins at 0.5 Å intervals, with an additional bin for distances exceeding 20 Å. The φ angle ranges from 0 to 180°, divided into 18 bins at 10° intervals, with an extra bin for non-contact scenarios. Both ω and θ range from -180 to 180°, segmented into 36 bins at 10° intervals, with an extra bin for non-contact scenarios. The ESM-2 features include a 1280-dimensional feature for each residue, containing their evolutionary information.

**Figure 1.**
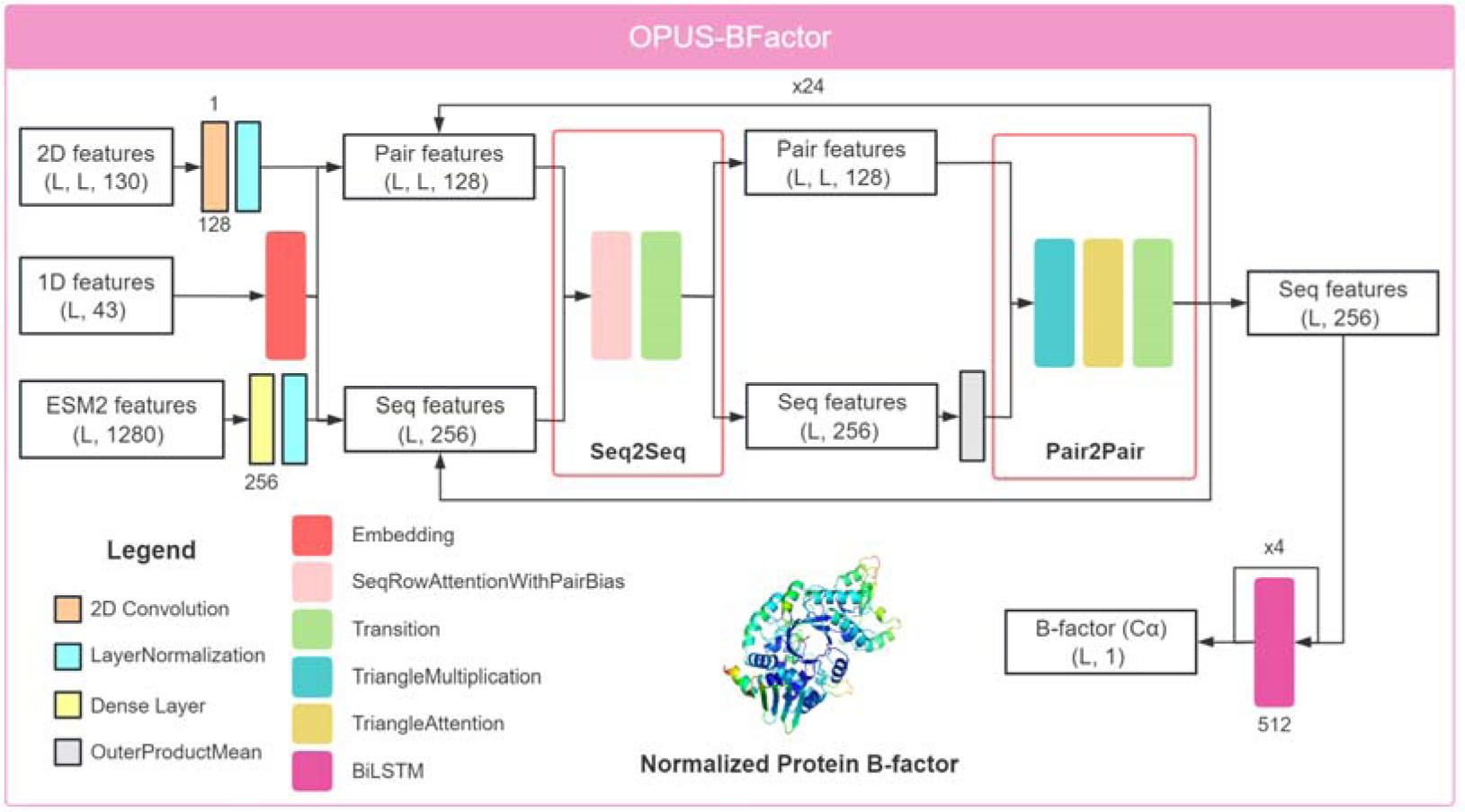
Overview of the OPUS-BFactor framework. OPUS-BFactor takes in three primary inputs: 1D protein sequence features, 2D residue-residue contact features, and 1D ESM-2 protein evolutionary features. In the first mode, OPUS-BFactor-seq, the structure-based features (first two) are set to zero, allowing predictions to be based solely on protein sequence. In the second mode, OPUS-BFactor-struct, the ESM-2 features are set to zero. For a target protein, OPUS-BFactor predicts the normalized B-factor of C_α_ atoms for all residues.

In OPUS-BFactor, the embedding module transforms the 1D features into sequence and pair features. The 2D features are processed through a 2D convolution layer and then added to the pair features, while the ESM-2 features are passed through a dense layer and added to the sequence features. Next, the RotaFormer module ^25^ is used to integrate the sequence and pair features. OPUS-BFactor employs 24 RotaFormers for feature extraction. After that, four BiLSTM layers ^23^ are used to further aggregate the sequence features and output the predicted B-factor value of C_α_ atoms. Following Pandey *et al.* ^12^, since the normalized B-factor has been shown to be more robust against experimental noise ^2^, we use the normalized B-factor in this study.

During training, we use the mean absolute error (MAE) loss between the predicted and actual normalized B-factor. We employ the Glorot uniform initializer and the Adam optimizer ^32^. The initial learning rate is set at 1e-3 and is halved after every two epochs. Training continues for a total of 6 epochs. OPUS-BFactor is developed using TensorFlow v2.4 ^33^ and trained on four NVIDIA Tesla V100 GPUs.

### Datasets

In OPUS-BFactor, we use the same training dataset as trRosetta ^30^, which includes 14,998 proteins after filtering the targets in which all residues have the same B-factor. To evaluate performance across various methods, we utilize three recently released test sets. The first, CAMEO65, was collected by Xu *et al.* ^34^, and contains 65 challenging targets released between May 2021 and October 2021 from the CAMEO website ^35^. After filtering, 62 targets remain. The second test set, CASP15, includes 44 targets available from the CASP website (http://predictioncenter.org). The third, CAMEO82, was collected by Xu *et al.* ^25^, and contains 82 targets released between May 2023 and August 2023 from the CAMEO website, with 75 targets remaining after filtering. In this study, we use the normalized B-factor (BC) for each C_α_ atom as the corresponding labels, calculated using the formula BC = (B - μ) / σ, where μ and σ are the mean and standard deviation of the unnormalized B-factor value (B) within the target protein.

### Performance Metrics

To evaluate the accuracy of each method, we use the average Pearson correlation coefficient (PCC) for each test set as our metric.

### Data and Software Availability

The code and pre-trained models of OPUS-BFactor as well as the datasets used in the study can be downloaded from http://github.com/OPUS-MaLab/opus_bfactor.

## Results

### Performance of different B-factor prediction methods

We evaluate the performance of OPUS-BFactor against a normal mode analysis (NMA)-based method, ProDy ^36^, and a deep learning-based method developed by Pandey *et al.* ^12^ across three test sets (CAMEO65, CASP15, and CAMEO82). As shown in Table 1, OPUS-BFactor consistently surpasses other methods in terms of average PCC on all three test sets. Additionally, the structure-based mode of OPUS-BFactor (OPUS-BFactor-struct) delivers better results than its sequence-based version (OPUS-BFactor-seq), indicating that structural information is crucial for accurate protein B-factor prediction. Figure 2 shows some prediction results from different methods.

**Figure 2.**
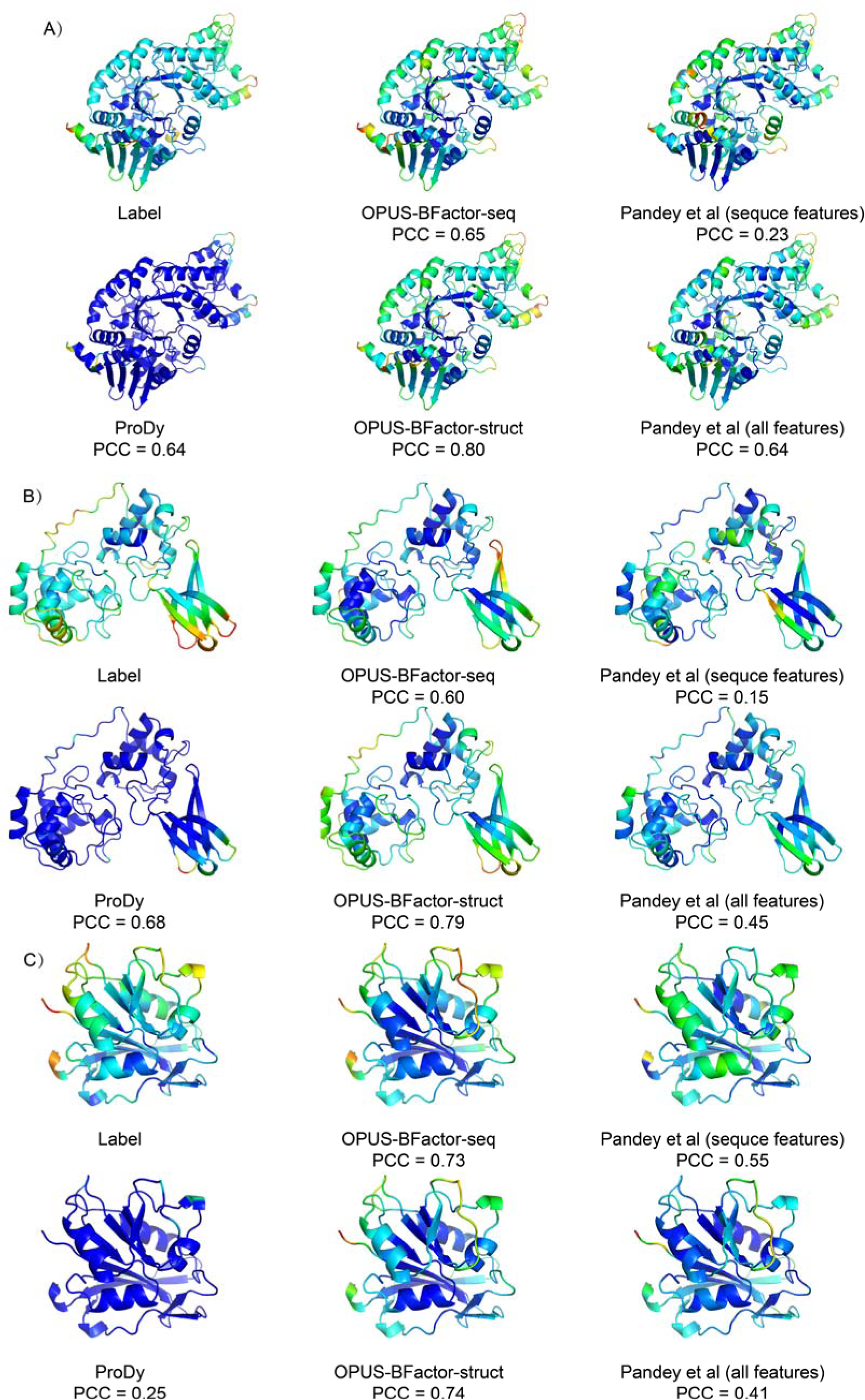
Protein B-factor prediction results of different methods. Results are colored in a spectrum (from red to blue) according to the B-factor of each C_α_ atom using PyMOL software. A) Prediction results on 2023-05-06_00000066_1. B) Prediction results on 2023-05-13_00000063_1. C) Prediction results on 2023-05-06_00000171_1.

**Table 1.**
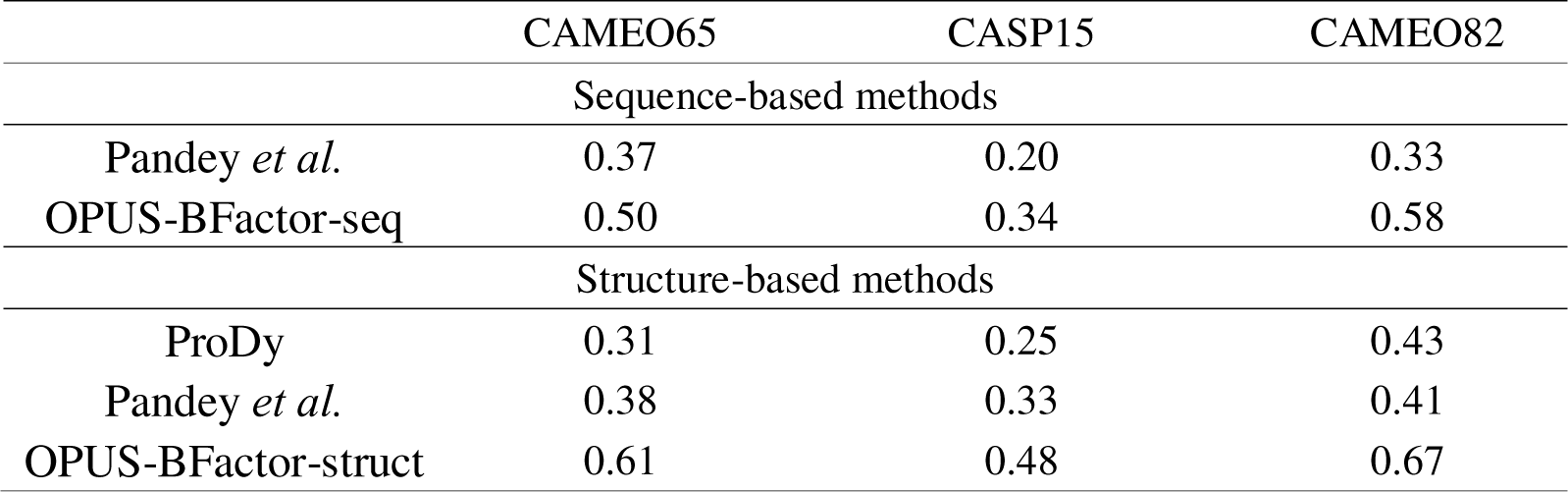
The average Pearson correlation coefficient (PCC) of different methods on each test set.

|  | CAMEO65 | CASP15 | CAMEO82 |
| --- | --- | --- | --- |
| Sequence-based methods |  |  |  |
| Pandey *et al.* | 0.37 | 0.20 | 0.33 |
| OPUS-BFactor-seq | 0.50 | 0.34 | 0.58 |
| Structure-based methods |  |  |  |
| ProDy | 0.31 | 0.25 | 0.43 |
| Pandey *et al.* | 0.38 | 0.33 | 0.41 |
| OPUS-BFactor-struct | 0.61 | 0.48 | 0.67 |

To investigate the correlation between protein B-factor and pLDDT from AlphaFold2 ^37^, we use AlphaFold2 to predict the 44 targets in the CASP15 test set and calculate the PCC between the real B-factor and pLDDT. Since the value of pLDDT represents the uncertainty of AlphaFold2’s prediction (with larger pLDDT values indicating less flexibility in the region), we use the negative pLDDT values for correlation calculation. The results show that the average PCC between the real B-factor and pLDDT is 0.23 on the CASP15 test set, which is lower than the average PCC of our sequence-based version OPUS-BFactor-seq (0.34), indicating a relatively low correlation between B-factor and pLDDT. This observation is consistent with the findings published by Carugo *et al*. ^38^. In Figure 3, we present some alignment results between the predicted values and real B-factors after normalization.

**Figure 3.**
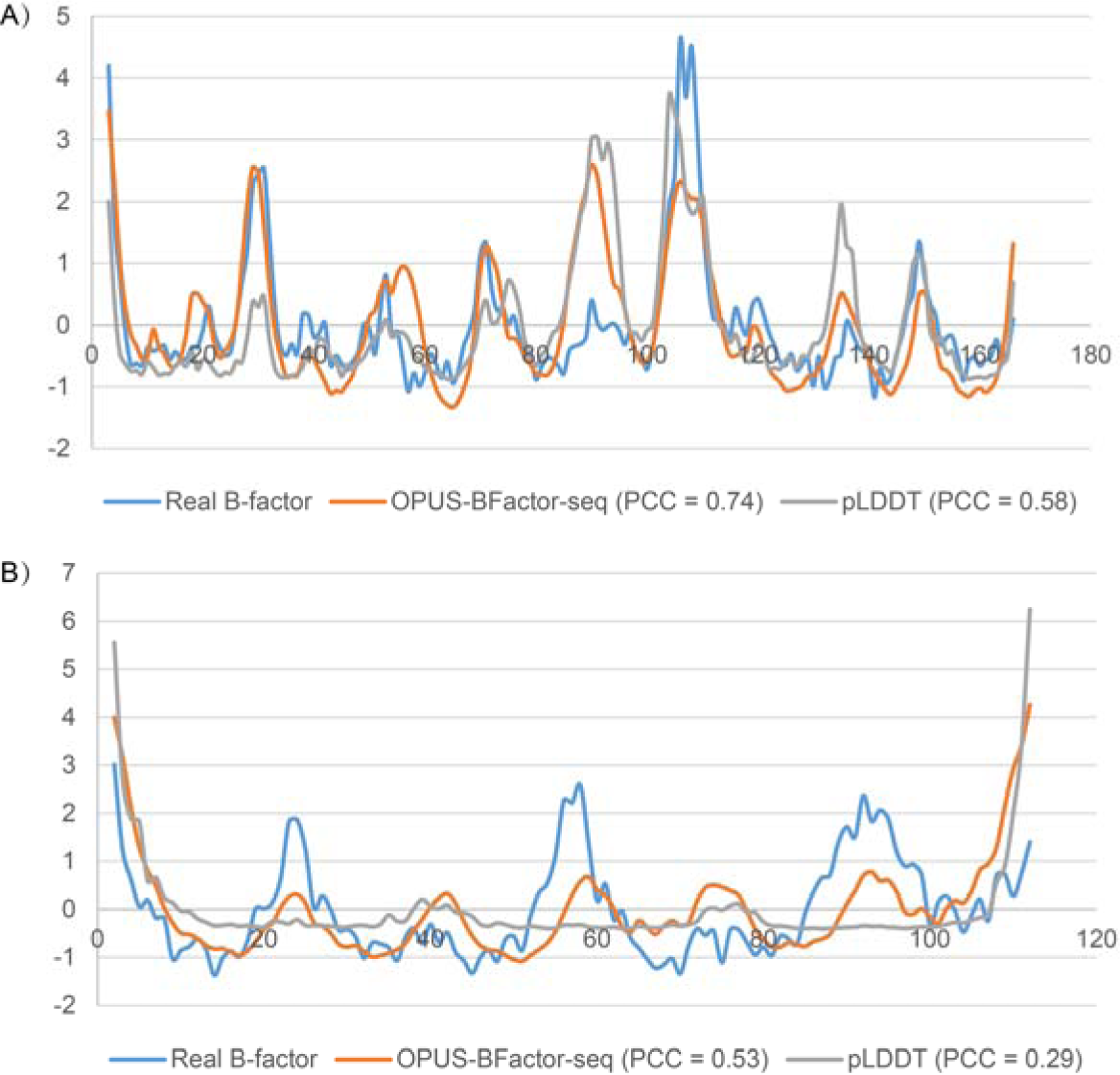
Alignment results between the real B-factor and the predicted B-factor from OPUS-BFactor-seq, as well as the pLDDT from AlphaFold2. All values are normalized in each protein. A) Results on T1187-D1. B) Results on T1106s2-D1.

### Ablation study of OPUS-BFactor

We evaluate the performance of sequence-based B-factor prediction models (OPUS-BFactor-seq) using different evolutionary profiles on the CAMEO82 test set. As shown in Figure 4, the model using ESM-2 features as inputs significantly improves prediction accuracy compared to models using one-hot encoding, HMM features, and PSSM features. This indicates that the performance of protein B-factor prediction can be enhanced by the utilization of more advanced evolutionary features.

**Figure 4.**
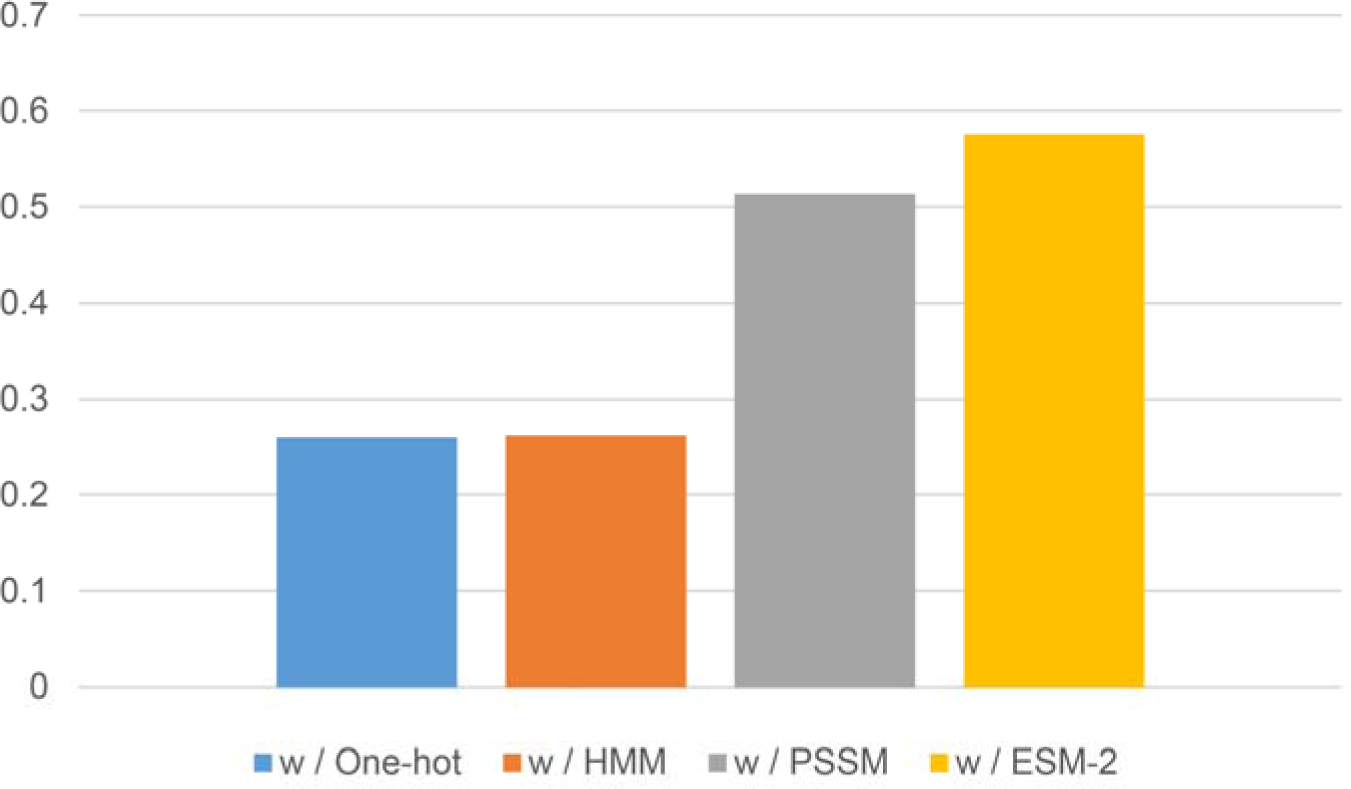
The average Pearson correlation coefficient (PCC) of OPUS-BFactor-seq using different evolutionary features as inputs on CAMEO82.

### Case study

In this study, we utilize OPUS-BFactor-seq to predict the B-factors for T4 lysozyme based on its sequence exclusively. As shown in Figure 5, we highlight two regions (Region A and B) in the prediction with relatively high values. The studies from other researchers show that Region A (D20-G23) corresponds to the active site of T4 lysozyme ^39^, and Region B (K35-L39) is a relatively flexible region as some researches indicate that an insertion or duplication of short peptide fragments in this area may cause a secondary structural transition (from helix to strand) ^40^.

**Figure 5.**
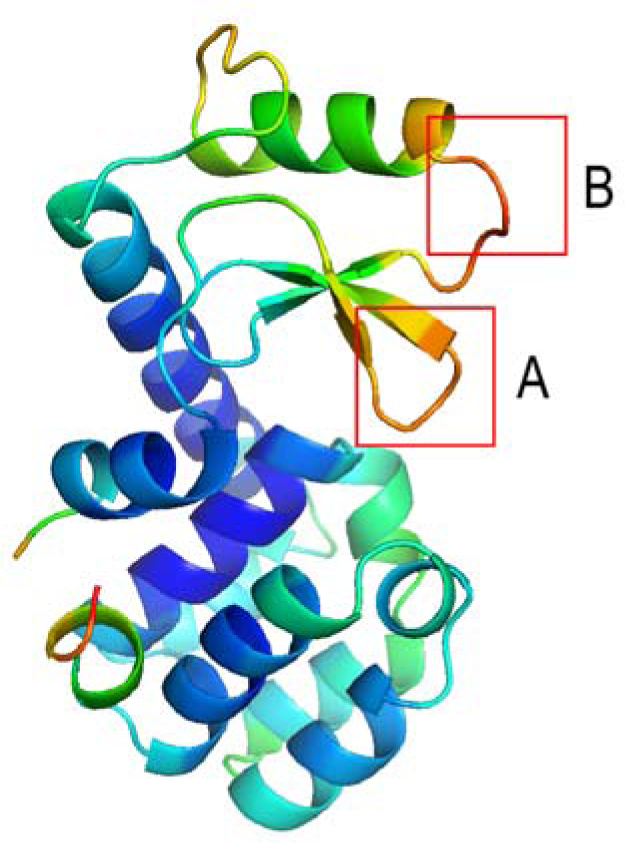
OPUS-BFactor-seq prediction results on T4 lysozyme using the sequence information exclusively. Results are colored in a spectrum (from red to blue) according to the B-factor of each C_α_ atom using PyMOL software. Region A includes residues between 20 and 23. Region B includes residues between 35 and 39.

## Concluding Discussion

In this study, we propose a protein B-factor prediction method called OPUS-BFactor, which operates in two modes: the first one (OPUS-BFactor-seq) uses sequence information exclusively, allowing predictions based solely on protein sequence; and the second one (OPUS-BFactor-struct) utilizes structural information, requiring the coordinates of backbone atoms in the target protein. The results (Table 1 and Figure 2) on three recently released test sets show that our method significantly outperforms other B-factor prediction methods. Meanwhile, the results highlight a performance gap between sequence-based and structure-based B-factor prediction models.

We also evaluate the correlation between real B-factors and predictions from OPUS-BFactor-seq, as well as the correlation between real B-factors and the pLDDT from AlphaFold2, both of which use sequence information exclusively. The results (Figure 3) show that OPUS-BFactor-seq delivers better results. Additionally, the results on T4 lysozyme (Figure 5) indicate that the regions with relatively high values of B-factors predicted by OPUS-BFactor-seq correspond with the active site and flexible regions of the target. Therefore, OPUS-BFactor-seq may serve as a useful tool for predicting protein properties related to the B-factor, such as flexibility, thermal stability, and regional activity.

Furthermore, the results (Figure 4) show that the performance of protein B-factor prediction may benefit from more advanced evolutionary features. In this case, protein B-factor prediction could serve as a valuable benchmark task for assessing protein language models. To facilitate this, we will make our formatted training and test sets, along with our code, available to all researchers.

## Acknowledgements

J. M. wants to thank the supports from National Key Research and Development Program of China (No. 2021YFF1200400), Science and Technology Innovation Plan Of Shanghai Science and Technology Commission (No. 23JS1400200), Shanghai Municipal Science and Technology Major Project (No.2018SHZDZX01), and ZJLab. G. X. wants to thank the support from National Natural Science Foundation of China (No. 32300535).

## Reference

1. Bramer, D.; Wei, G. W., Blind prediction of protein B-factor and flexibility. J Chem Phys 2018, 149 (13).

2. Sun, Z. T.; Liu, Q.; Qu, G.; Feng, Y.; Reetz, M. T., Utility of B-Factors in Protein Science: Interpreting Rigidity, Flexibility, and Internal Motion and Engineering Thermostability. Chem Rev 2019, 119 (3), 1626–1665.

3. Vihinen, M.; Torkkila, E.; Riikonen, P., Accuracy of Protein Flexibility Predictions. Proteins 1994, 19 (2), 141–149.

4. Karplus, P. A.; Schulz, G. E., Prediction of Chain Flexibility in Proteins - a Tool for the Selection of Peptide Antigens. Naturwissenschaften 1985, 72 (4), 212–213.

5. Parthasarathy, S.; Murthy, M. R. N., Protein thermal stability:: insights from atomic displacement parameters (values). Protein Eng 2000, 13 (1), 9–13.

6. Yuan, Z.; Zhao, J.; Wang, Z. X., Flexibility analysis of enzyme active sites by crystallographic temperature factors. Protein Eng 2003, 16 (2), 109–114.

7. Radivojac, P.; Obradovic, Z.; Smith, D. K.; Zhu, G.; Vucetic, S.; Brown, C. J.; Lawson, J. D.; Dunker, A. K., Protein flexibility and intrinsic disorder. Protein Sci 2004, 13 (1), 71–80.

8. Atilgan, A. R.; Durell, S. R.; Jernigan, R. L.; Demirel, M. C.; Keskin, O.; Bahar, I., Anisotropy of fluctuation dynamics of proteins with an elastic network model. Biophys J 2001, 80 (1), 505–515.

9. Ma, J. P., Usefulness and limitations of normal mode analysis in modeling dynamics of biomolecular complexes. Structure 2005, 13 (3), 373–380.

10. Pan, X. Y.; Shen, H. B., Prediction of Protein B-factor Profile based on Feature Selection and Kernel Learning. Proceedings of the 2009 Chinese Conference on Pattern Recognition and the First Cjk Joint Workshop on Pattern Recognition, Vols 1 and 2 2009, 588-592.

11. Yuan, Z.; Bailey, T. L.; Teasdale, R. D., Prediction of protein B-factor profiles. Proteins 2005, 58 (4), 905–912.

12. Pandey, A.; Liu, E.; Graham, J.; Chen, W.; Keten, S., B-factor prediction in proteins using a sequence-based deep learning model. Patterns 2023, 4 (9).

13. Pang, Y.-P., Use of multiple picosecond high-mass molecular dynamics simulations to predict crystallographic B-factors of folded globular proteins. Heliyon 2016, 2 (9).

14. Wang, Q.; Xiao, X.; Miao, Z.; Zhang, X.; Jiang, B.; Liu, M., Prediction of Protein B-factor Profiles based on Bidirectional Long Short-Term Memory Network. ChemRxiv 2023.

15. Lu, M. Y.; Ming, D. M.; Ma, J. P., fSUB: Normal Mode Analysis with Flexible Substructures. J Phys Chem B 2012, 116 (29), 8636–8645.

16. Poon, B. K.; Chen, X. R.; Lu, M. Y.; Vyas, N. K.; Quiocho, F. A.; Wang, Q. H.; Ma, J. P., Normal mode refinement of anisotropic thermal parameters for a supramolecular complex at 3.42-A crystallographic resolution. P Natl Acad Sci USA 2007, 104 (19), 7869–7874.

17. Haliloglu, T.; Bahar, I.; Erman, B., Gaussian dynamics of folded proteins. Phys Rev Lett 1997, 79 (16), 3090–3093.

18. Bahar, I.; Atilgan, A. R.; Erman, B., Direct evaluation of thermal fluctuations in proteins using a single-parameter harmonic potential. Fold Des 1997, 2 (3), 173–181.

19. Zhang, H.; Jiang, T.; Shan, G. G.; Xu, S. Q.; Song, Y. J., Gaussian network model can be enhanced by combining solvent accessibility in proteins. Sci Rep-Uk 2017, 7.

20. Yang, J. Y.; Wang, Y.; Zhang, Y., ResQ: An Approach to Unified Estimation of -Factor and Residue-Specific Error in Protein Structure Prediction. J Mol Biol 2016, 428 (4), 693–701.

21. Bramer, D.; Wei, G. W., Multiscale weighted colored graphs for protein flexibility and rigidity analysis. J Chem Phys 2018, 148 (5).

22. LeCun, Y.; Bengio, Y.; Hinton, G., Deep learning. Nature 2015, 521 (7553), 436–444.

23. Hochreiter, S.; Schmidhuber, J., Long short-term memory. Neural Comput 1997, 9 (8), 1735–80.

24. Lin, Z. M.; Akin, H.; Rao, R. S.; Hie, B.; Zhu, Z. K.; Lu, W. T.; Smetanin, N.; Verkuil, R.; Kabeli, O.; Shmueli, Y.; Costa, A. D.; Fazel-Zarandi, M.; Sercu, T.; Candido, S.; Rives, A., Evolutionary-scale prediction of atomic-level protein structure with a language model. Science 2023, 379 (6637), 1123–1130.

25. Xu, G.; Luo, Z.; Yan, Y.; Wang, Q.; Ma, J., OPUS-Rota5: A highly accurate protein side-chain modeling method with 3D-Unet and RotaFormer. Structure 2024, 32 (7), 1001–1010.e2.

26. Hanson, J.; Paliwal, K.; Litfin, T.; Yang, Y. D.; Zhou, Y. Q., Improving prediction of protein secondary structure, backbone angles, solvent accessibility and contact numbers by using predicted contact maps and an ensemble of recurrent and residual convolutional neural networks. Bioinformatics 2019, 35 (14), 2403–2410.

27. Xu, G.; Wang, Q. H.; Ma, J. P., OPUS-TASS: a protein backbone torsion angles and secondary structure predictor based on ensemble neural networks. Bioinformatics 2020, 36 (20), 5021–5026.

28. Lu, M. Y.; Dousis, A. D.; Ma, J. P., OPUS-PSP: An orientation-dependent statistical all-atom potential derived from side-chain packing. J Mol Biol 2008, 376 (1), 288–301.

29. Xu, G.; Ma, T. Q.; Zang, T. W.; Sun, W. T.; Wang, Q. H.; Ma, J. P., OPUS-DOSP: A Distance- and Orientation-Dependent All-Atom Potential Derived from Side-Chain Packing. J Mol Biol 2017, 429 (20), 3113–3120.

30. Yang, J. Y.; Anishchenko, I.; Park, H.; Peng, Z. L.; Ovchinnikov, S.; Baker, D., Improved protein structure prediction using predicted interresidue orientations. P Natl Acad Sci USA 2020, 117 (3), 1496–1503.

31. Xu, G.; Wang, Q. H.; Ma, J. P., OPUS-X: an open-source toolkit for protein torsion angles, secondary structure, solvent accessibility, contact map predictions and 3D folding. Bioinformatics 2022, 38 (1), 108–114.

32. Kingma, D. P.; Ba, J., Adam: A Method for Stochastic Optimization. Proceedings of the 3rd International Conference on Learning Representations 2015.

33. Abadi, M.; Barham, P.; Chen, J. M.; Chen, Z. F.; Davis, A.; Dean, J.; Devin, M.; Ghemawat, S.; Irving, G.; Isard, M.; Kudlur, M.; Levenberg, J.; Monga, R.; Moore, S.; Murray, D. G.; Steiner, B.; Tucker, P.; Vasudevan, V.; Warden, P.; Wicke, M.; Yu, Y.; Zheng, X. Q., TensorFlow: A system for large-scale machine learning. Proceedings of the 12th USENIX Symposium on Operating Systems Design and Implementation 2016, 265-283.

34. Xu, G.; Wang, Q. H.; Ma, J. P., OPUS-Mut: Studying the Effect of Protein Mutation through Side-Chain Modeling. J Chem Theory Comput 2023, 19 (5), 1629–1640.

35. Haas, J.; Barbato, A.; Behringer, D.; Studer, G.; Roth, S.; Bertoni, M.; Mostaguir, K.; Gumienny, R.; Schwede, T., Continuous Automated Model EvaluatiOn (CAMEO) complementing the critical assessment of structure prediction in CASP12. Proteins 2018, 86, 387–398.

36. Bakan, A.; Meireles, L. M.; Bahar, I.,: Protein Dynamics Inferred from Theory and Experiments. Bioinformatics 2011, 27 (11), 1575–1577.

37. Jumper, J.; Evans, R.; Pritzel, A.; Green, T.; Figurnov, M.; Ronneberger, O.; Tunyasuvunakool, K.; Bates, R.; Zidek, A.; Potapenko, A.; Bridgland, A.; Meyer, C.; Kohl, S. A. A.; Ballard, A. J.; Cowie, A.; Romera-Paredes, B.; Nikolov, S.; Jain, R.; Adler, J.; Back, T.; Petersen, S.; Reiman, D.; Clancy, E.; Zielinski, M.; Steinegger, M.; Pacholska, M.; Berghammer, T.; Bodenstein, S.; Silver, D.; Vinyals, O.; Senior, A. W.; Kavukcuoglu, K.; Kohli, P.; Hassabis, D., Highly accurate protein structure prediction with AlphaFold. Nature 2021, 596 (7873), 583-+.

38. Carugo, O., pLDDT Values in AlphaFold2 Protein Models Are Unrelated to Globular Protein Local Flexibility. Crystals 2023, 13 (11).

39. Weaver, L. H.; Matthews, B. W., Structure of Bacteriophage-T4 Lysozyme Refined at 1.7 a Resolution. J Mol Biol 1987, 193 (1), 189–199.

40. Kaur, H.; Sasidhar, Y. U., Molecular dynamics study of an insertion/duplication mutant of bacteriophage T4 lysozyme reveals the nature of α → β transition in full protein context. Phys Chem Chem Phys 2013, 15 (20), 7819–7830.

